# Extensive Chromatin Structure-Function Association Revealed by Accurate Compartmentalization Characterization

**DOI:** 10.1101/2021.09.17.460762

**Authors:** Zi Wen, Weihan Zhang, Quan Zhong, Jinsheng Xu, Chunhui Hou, Zhaohui Qin, Li Li

## Abstract

Chromosome conformation capture-based experiments have shown that eukaryotic chromosomes are partitioned into A and B compartments conventionally identified by the first eigenvector (EV1) of dimension reduction methods. However, many genomic regions show marginal EV1 values, indicating the ambiguity of A/B compartment scheme on these regions. We develop MOSAIC (MOdularity and Singular vAlue decomposition-based Identification of Compartments), an accurate compartmental state detection scheme. MOSAIC reveals that those ambiguous regions segregate into two additional compartmental states, which typically correspond to small genomic regions flanked by large canonical A/B compartments with opposite activities. They are denoted as micro-compartments accordingly. In contrast to the canonical A/B compartments, micro-compartments cover ~30% of the genome and are highly dynamic between cell types. More importantly, distinguishing the micro-compartments underpins accurate characterization of chromatin structure-function relationship. By applying MOSAIC to GM12878 and K562 cells, we identify *CD86*, *ILDR1* and *GATA2* which show concordance between gene expression and compartmental states beyond the scheme of A/B compartments. Taken together, MOSAIC uncovers fine-scale and dynamic structures underlying canonical A/B compartments. Our results suggest dynamic chromatin compartmentalization is underlying transcriptional regulation and disease.

## Introduction

Modularity, a widely recognized principle of living systems, has been observed in many aspects of biological organization (Wagner et al. 2007). In the eukaryotic genome, the highest level of physical modularity is the separation of genetic material into chromosomes that occupy discrete territories inside the nucleus. Chromosome conformation capture (3C)-based high-throughput technologies have revealed rich modular features in 3D chromatin architecture at fine scales (Gibcus and Dekker 2013; Schoenfelder and Fraser 2019). Plaid patterns were observed at the megabase scale in intra-chromosomal heatmaps generated by the first high-throughput chromosome conformation capture (Hi-C) experiments (Lieberman-Aiden et al. 2009) on human cells. These patterns suggested the existence of two segregated groups of chromatin in which regions within the same group are in closer spatial proximity than regions across groups, a hallmark of modularity. The two groups, or structural modules, were named the A and B compartments, corresponding to transcriptionally active and silent chromatin, respectively. Principal component analysis (PCA) of the correlation heatmap yields the first eigenvector (EV1), whose sign designates whether a region is in compartment A or B. Due to its simplicity in implementation, straightforwardness in interpretation, and robustness to noise, eigenvector decomposition methods, including PCA and singular value decomposition (SVD), are the standard procedures for compartmentalization analysis (Schmitt et al. 2016b; Durand et al. 2016; Wolff et al. 2018).

Compartmental organization of chromosomes has been confirmed in other animals (Dixon et al. 2012; Rowley et al. 2017), plants (Dong et al. 2017), and archaea (Takemata et al. 2019). Although it disappears temporarily in certain stages of mitosis (Naumova et al. 2013) and zygotic genome activation (Du et al. 2017; Ke et al. 2017; Niu et al. 2021), chromosome compartmentalization is a widespread pattern in a variety of cell types and biological processes. Given the mechanistic connection to phase separation (Larson et al. 2017; Strom et al. 2017) and the distinction in gene activity between compartments, the role of compartmentalization in chromatin functions, especially transcriptional regulation, becomes an increasingly important question (Hildebrand and Dekker 2020).

While assignment of A/B compartments by EV1 can capture the overall plaid pattern of individual intra-chromosomal heatmaps, studies based on trans contracts have further divided the human genome into six subcompartments (Rao et al. 2014; Ashoor et al. 2020; Xiong and Ma 2019). Recently, a computational framework that integrates TSA-seq, DamID and Hi-C data provides spatial interpretation for Hi-C subcompartments relative to multiple nuclear compartments (Wang et al. 2021). Another method starts with compartmental domain identification, followed by hierarchical clustering on the domains, then obtains eight subcompartment types (Liu et al. 2021). These results suggest that simply partitioning chromatin into two compartments is insufficient to capture the full complexity of chromatin compartmentalization. In practice, however, when bipartite compartment analysis and subcompartment analysis are compared, the former is still more widely applied (Furlan-Magaril et al. 2021; Belaghzal et al. 2021; Sati et al. 2020) considering its straightforward connection to chromatin activity and robustness to noise (Schmitt et al. 2016b). Thus, it is important to design a method which advance the eigenvector decomposition to more refined levels while keeping its advantages over the subcompartment methods.

In this study, we showed that modularity can be used to quantify chromatin compartmentalization. Based on this observation, we developed a modularity-based compartmentalization analysis framework named MOSAIC (MOdularity and Singular vAlue decomposition-based Identification of Compartments), using intra-chromosomal contact information from Hi-C. After performing dimension reduction by SVD, we systematically evaluated the structural and functional implications of additional EVs orthogonal to EV1. While EV1 showed the strongest correlation with active histone marks as known, we found that the second EV (EV2) strongly correlated with H3K27me3, highlighting facultative heterochromatin, which typically consists of small regions interspersed within large areas of the A compartment. Totally, ~ 30% of the genome do not belong to canonical A/B compartments. These initially ambiguous regions harboring marginal EV1 values can be clarified and segregated into two additional components, which we termed micro-compartments given their smaller size and genomic neighborhood with respect to canonical A/B compartments. Through the top EVs, MOSAIC intuitively connected 3D chromatin architecture to 1D chromatin states and activities, including histone modifications and transcription. More importantly, when applying MOSAIC to GM12878 and K562 cells, we identified much more genes that exhibit concordance between expression and compartmental states when compared to results obtained using conventional A/B compartments. Furthermore, the case studies on *HoxA* gene cluster and *GATA2* exemplify the power of MOSAIC in revealing such structurally delicate but biologically essential features, and the limitations of the A/B compartment scheme.

## Results

### MOSAIC overview and EV exploration

To identify compartmental components in an individual chromosome, we started with the 100 kb-binned intra-chromosomal “observed over expected” (O/E) matrix (Lieberman-Aiden et al. 2009) and used Hi-C data from the GM12878 cell line (Rao et al. 2014). First, we applied SVD to the O/E matrix (Fig. 1A). For some chromosomes, one of the top PCs reflects the long and short arms (Schmitt et al. 2016b), potentially due to the barrier role of centromeres in suppressing contacts across chromosome arms (Muller et al. 2019). To disentangle the impact of the centromere effect on compartmentalization analysis, we quantified EVs in terms of arm bias, and the O/E matrices showing a strong centromere effect were corrected accordingly (see Methods for details).

**Figure 1.**
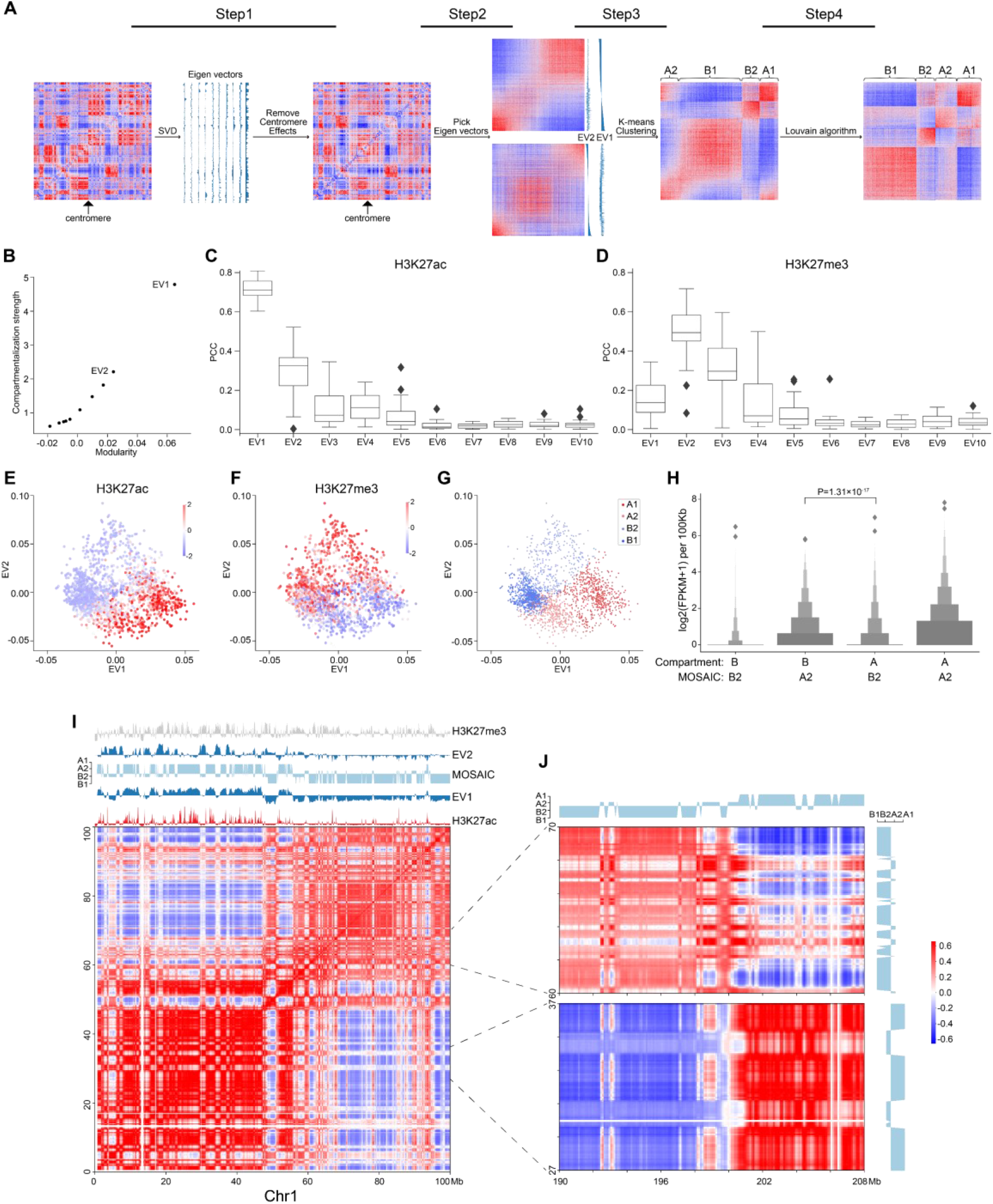
Workflow of MOSAIC and overview of compartmental states. (*A*) MOSAIC workflow starts with an individual intra-chromatin O/E matrix. After two rounds of SVD, potential centromere effect is removed from the matrix, and the two eigenvectors that can best reflect the structural features are selected for K-means clustering. Finally, a modified Louvain algorithm is applied to refine the results of K-means clustering. (*B*) The average compartmentalization strength of all chromosomes versus the average modularity of all chromosomes for the top-ten EVs in terms of modularity for each chromosome. (*C-D*) Pearson correlation coefficients between the top-ten EVs sorted by modularity and H3K27ac (*C*) and H3K27me3 (*D*), respectively. (*E-F*) Enrichment of H3K27ac (*E*) and H3K27me3 (*F*) for all 100 kb bins of chromosome 1 in relation to EV1 and EV2, respectively. The enrichment values are ZSCOR-normalized. (*G*) Compartmental states of all 100 kb bins of chromosome 1 in relation to EV1 and EV2. (*H*) Gene expression for the A2 and B2 regions identified by the MOSAIC and A/B compartment schemes (P value calculated by one-tailed Mann– Whitney rank test). (*I*) Hi-C correlation heatmap of Chr1: 0–100 Mb along with EV1, EV2, H3K27me3, H3K27ac, and compartmental states. (*J*) Hi-C correlation heatmap of Chr1: 27–37 Mb, 60–70 Mb and the distal region Chr1: 190–208 Mb.

With the O/E matrices corrected for centromere effect, we performed another round of SVD and explored the structural and functional implications of the top EVs. From the perspective of modularity, compartmentalization is a modular feature, with the A/B compartments representing two structural modules of the genome. To quantify an EV in terms of structural segregation, a modularity score (Newman 2006), termed *Q*_*EV*_, can be assigned, supposing a bipartition based on its sign (Newman 2013) along the chromosome. The top-ten EVs were re-ranked by their *Q*_*EV*_ in descending order for each chromosome. EV1, which corresponds to the A/B compartments, gave the highest *Q*_*EV*_ (Supplemental Fig. S1A). We also adapted compartmentalization strength as previously described (Schwarzer et al. 2017) to quantify the structural segregation represented by each eigenvector, and found that the compartmentalization strength was quantitatively consistent with modularity (Fig. 1B; Supplemental Fig. S1B). As expected, EV1 possesses both the highest compartmentalization and the highest *Q*_*EV*_, followed by EV2. Therefore, MOSAIC adopts modularity as the target function for compartment identification.

To quantify the epigenomic and functional implications of EVs, their correlation to various histone modifications was calculated and denoted as *R*_*EV*_. Due to arbitrary sign designation, we used the absolute value to measure the correlation strength. For the top-ten EVs, *Q*_*EV*_ and *R*_*EV*_ were highly correlated (Fig. 1C; Supplemental Fig. S1C-D), indicating concordance between chromatin structure and epigenetic states. In other words, EVs with high modularity were also highly correlated to histone modification signals. Such a monotonic relationship held for all histone modifications, with H3K27me3 to be the only exception (Fig. 1D; Supplemental Fig. S1C); for H3K27me3, the EV with the highest correlation was EV2 rather than EV1 (Fig. 1D; Supplemental Fig. S1D). In contrast to the marginal correlation between EV1 and H3K27me3, EV2 and H3K27me3 had a Pearson correlation coefficient (PCC) around 0.5. This prominent distinction prompted us to interrogate EV2 and its relation to H3K27me3.

To further explore the relationship between EVs and histone modifications, we made scatter plots using EV1 and EV2, and examined the patterns of histone modifications of all 100 kb-binned regions in chromosome 1 (Fig. 1E–F). For H3K27ac (Fig. 1E), as expected, the regions with the most enriched and depleted signals lay within the two extremes along EV1, corresponding to the typical A and B compartments, respectively. For H3K27me3, which lacked a clear level gradient along EV1, EV2 became a strong indicator (Fig. 1F). Moreover, the span of EV2 was largely orthogonal to EV1, as indicated by the convex outline of the data points. According to the algebraic interpretation of EVs, the magnitude of EV for a region represents the strength of the noted pattern. As shown in Figure 1E and 1F, regions with extreme EV2 values also have marginal EV1, indicating that the regions ambiguous under EV1 are in fact structurally distinctive, as pinpointed by EV2 (Supplemental Fig. S1E-F). Taken together, these findings indicate that EV1 and EV2 are complementary in characterizing the structural and functional states of chromatin.

Based on above results, EV1 and EV2 were most representative and informative in terms of both structural modularity and epigenomic relatedness. Therefore, we chose them to conduct K-means clustering to identify compartmental states. We employed the sum of squared errors (SSE) to determine the optimal number of clusters (Supplemental Fig. S1G). Accordingly, cluster number of four were adopted in subsequent analysis. Because compartmentalization is a global feature, we sought to optimize the identification of compartmental states by Louvain algorithm (Blondel et al. 2008) to maximize modularity and obtained the final four compartmental states (Fig. 1A; see Methods for details).

### MOSAIC captures structural and functional features more accurately than A/B compartments

The distribution of the four modules for chromosome 1 is shown in Figure 1G. The two modules lying to the left and right, well separated by EV1, correspond to canonical A/B compartments, and were therefore denoted as A1 and B1 (Supplemental Fig. S1H). The other two modules on the top and bottom were mostly ambiguous in the A/B compartment scenario. After expanding to the 2D space of EV1 and EV2, they are clearly segregated along the dimension of EV2 and coincide with by H3K27me3 levels. Because of the repressive nature of H3K27me3, the modules with and without H3K27me3 signals were denoted as B2 and A2, respectively. To further confirm the activity distinction of A2 and B2, we checked the discrepant regions between our partition and the A/B partition based on only the sign of EV1 (Supplemental Fig. S1H). In the matching matrix, the regions originally considered B, but now identified as A2, are transcriptionally more active than the regions originally considered A but now identified as B2 (Fig. 1H), justifying the discriminating power of EV2 in these regions ambiguous under EV1. In summary, each individual chromosome of the genome was partitioned into four compartmental states, labeled A1, B1, A2, and B2 (Supplemental Fig. S2A).

To examine our partition results, we visually inspected intra-chromosomal heatmaps along with compartmental states identified by MOSAIC. As an example, Figure 1I shows the correlation heatmap of the initial 100 Mb of chromosome 1. This area can be coarsely split from near the center, as noted by the two large predominantly red squares along the diagonal of the heatmap. Default A/B compartments marked by the sign of EV1 correctly reflect this coarse-grained pattern, with the two halves of EV1 profile having opposite signs (Fig. 1I). Nevertheless, interspersed on each half, we can see plentiful secondary and fine plaid patterns on the heatmap. These patterns are visualized as fine bands in alternating white and red that are lighter than the surrounding background of the large red squares. Notably, on the EV1 profile, most of these bands just correspond to local depressions without sign switching. That is, most of these fine patterns are not identified as compartment-switching between conventional A and B compartment. Strikingly, such moderate contrasts, although secondary to the distinction of A/B compartments, are highlighted by EV2 with sharp peaks or plateaus on the positive or negative side. Furthermore, the plaid pattern extends and spans the whole chromosome, including distal regions on the other arm (Fig. 1J), confirming the global compartmental features of the newly identified modules. Such patterns of the interspersed regions, which also correspond to the points on the top and bottom areas of Figure 1G, ground the structural significance of EV2 and the rationale of distinguishing these regions from the surrounding typical A/B compartments.

Consistent with their strong correlation with H3K27me3 (Fig. 1D), the EV2 peaks were specifically marked by H3K27me3 signals (Fig. 1F). Such coincidence points to an intuitive interpretation of EV2 and supports the heterochromatic nature of the corresponding regions. Therefore, these regions were denoted as B2 to differentiate them from B1, which represents constitutive heterochromatin. Taken together, the area on chromosome 1 confirms the orthogonality of EV1 and EV2 in capturing the patterns in heatmaps, and supports the refined partition of the chromosome into four compartmental states, with the newly identified A2 and B2 which are typically small regions surrounded by large areas of B1 and A1, respectively.

### Whole genome characterization of micro-compartment and compartment borders

To examine our findings on all chromosomes, we conducted statistical analyses on the whole genome. The newly identified A2 and B2 regions exhibit features distinct from those of A1 and B1, which correspond to the conventional A and B compartments. In terms of genomic constitutions, A1 and B1 sum to 70.9% of the genome. A2 and B2 constitute the other 29.1% (Fig. 2A). Nevertheless, the number of regions in each compartmental state is comparable (Supplemental Fig. S2B), while their length distributions differ dramatically (Fig. 2B). Consistent with the example shown in Figure 1I, regions of A2 and B2 are significantly shorter than those of A1 and B1. Therefore, we denoted A2 and B2 as micro-compartment.

**Figure 2.**
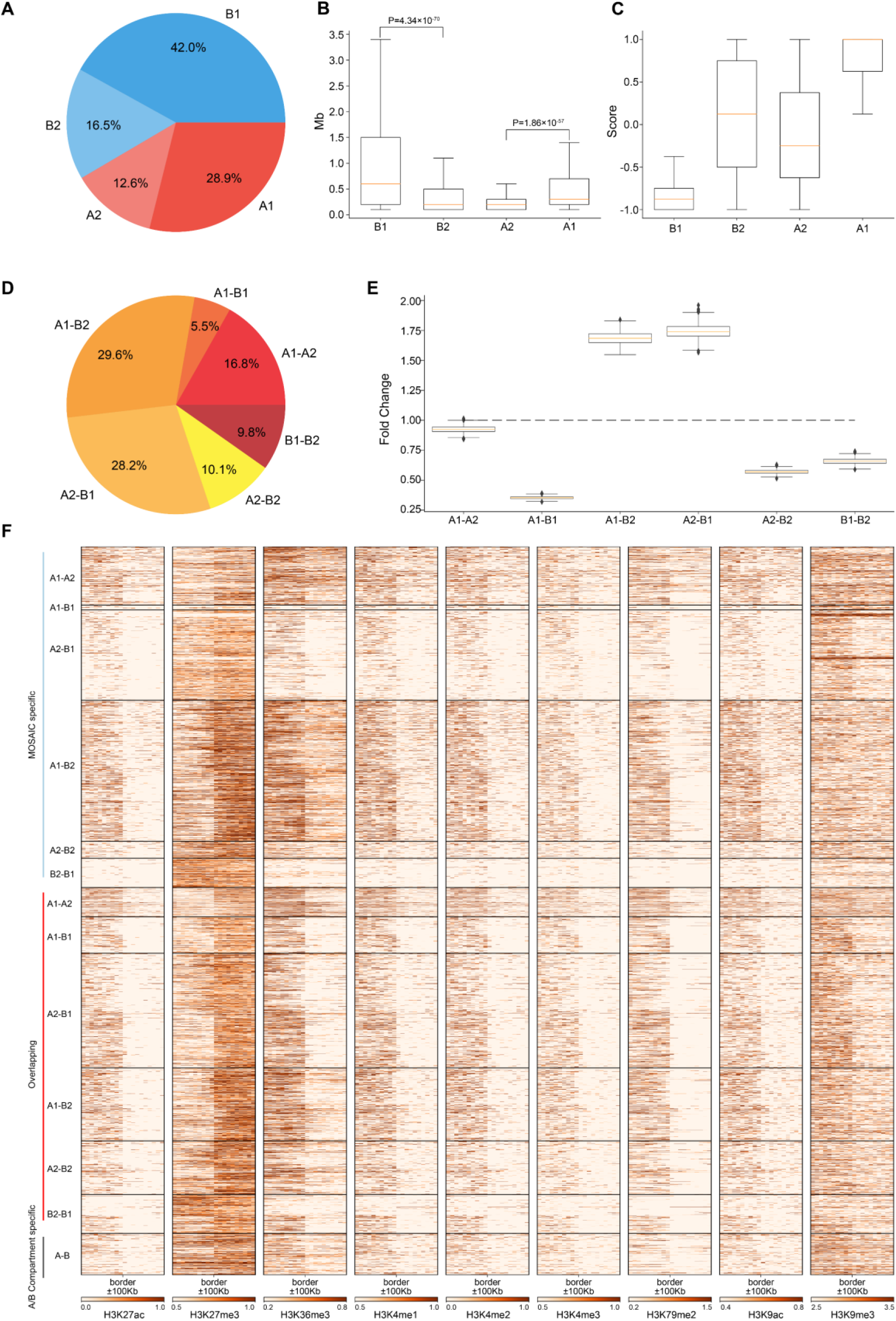
Regions of A2 and B2 are short, constitute a minor proportion of the genome and are embedded within B1 and A1, respectively. (*A*) Proportions of four compartmental states throughout the genome. (*B*) Length distributions of four compartmental states throughout the genome (P value calculated by one-tailed Mann–Whitney rank test). (*C*) Compartmental conservations of four compartmental states in 16 cell lines. The values range from −1 to 1. A value of 1 means that the region is in the A compartment in all 16 cell types; a value of −1 means that it is in the B compartment in all 16 cell types; and a value of 0 means that it is in A compartment in 8 cell lines and in B compartment in the other 8 cell lines. (*D*) Proportions of border types in terms of neighboring compartmental states. (*E*) Distribution of enrichment of true border types relative to random border types with neighborhoods shuffled 1,000 times. (*F*) Heatmaps of various histone modifications at the grouped border types.

To characterize the degree of compartmental conservation, we examined the results of A/B compartmentalization in 16 cell lines (Schmitt et al. 2016a) and defined a compartmental conservation score for each region by counting the A/B compartment assignment in the cell lines. As shown in Figure 2C, A1 and B1 correspond to consensus A/B compartments. By contrast, A2 and B2 tend to switch between A and B compartments, thus correspond to dynamic and cell-type specific compartmental regions from the perspective of A/B compartments.

Given the four compartmental states, there will be six combinations in terms of genomic neighborhood. Among the six neighboring types, majority are A1-B2 and A2-B1 (Fig. 2D). These two types are highly enriched relative to the results of compartment identity permutations, whereas the other four neighboring types are under-represented (Fig. 2E). Combined with the region length distributions (Fig. 2B), these features coincide with the observations on chromosome 1 that B2 and A2 are embedded within large areas of A1 and B1, respectively. The depletion of A1-A2 and B1-B2 means that subtypes of euchromatin/heterochromatin do not tend to be next to each other. Notably, the depletion of A1-B1 suggests that drastic switches of compartmental states are rarer than expected. Instead, genomic neighborhood favors moderate switches between euchromatin heterochromatin, e.g., A1-B2 and A2-B1.

Analysis of the compartment borders revealed 2,205 MOSAIC-specific borders, 270 A/B compartment-specific borders and 1,849 overlapping borders (Supplemental Fig. S2C). 99.2% of MOSAIC-specific borders were associated with A2/B2 (Fig. 2F), confirming that the MOSAIC-specific borders are the consequence of the newly identified micro-compartments. We found that the MOSAIC-specific borders have distinct patterns of histone modification transition depending on the border types (Fig. 2F). For example, the transition patterns of H3K27me3 and H3K36me3 at the A1-B2 borders are different from those at the A2-B1 borders. For the overlapping borders, MOSAIC revealed that they can be distinguished and divided into six types with distinct epigenetic patterns, which the A/B compartment scheme cannot discern. Together, these results tell us that micro-compartments are fragmented regions embedded in large areas of A1 and B1. These architectural features are reinforced by the distinct histone modification transitions at corresponding compartment borders.

### Compartmental states show distinct epigenetic and functional features

Our analysis of Hi-C data revealed that individual chromosomes segregate into four compartmental components. To characterize their epigenetic and functional features, we explored the pattern of histone modifications and replication timing using a previously described method (Rao et al. 2014) (Fig. 3A). The results revealed that A1 and A2 regions are both enriched in histone modifications representing euchromatin or transcriptional activities, with A1 enriched to a higher degree than A2. B2 regions are specifically enriched in H3K27me3 signals. B1 regions are barely enriched in any histone modifications, indicative of constitutive heterochromatin. Consistent with this, the replication timing enrichment of S3, S4, and G2 in these regions reveals that they are late replicating.

**Figure 3.**
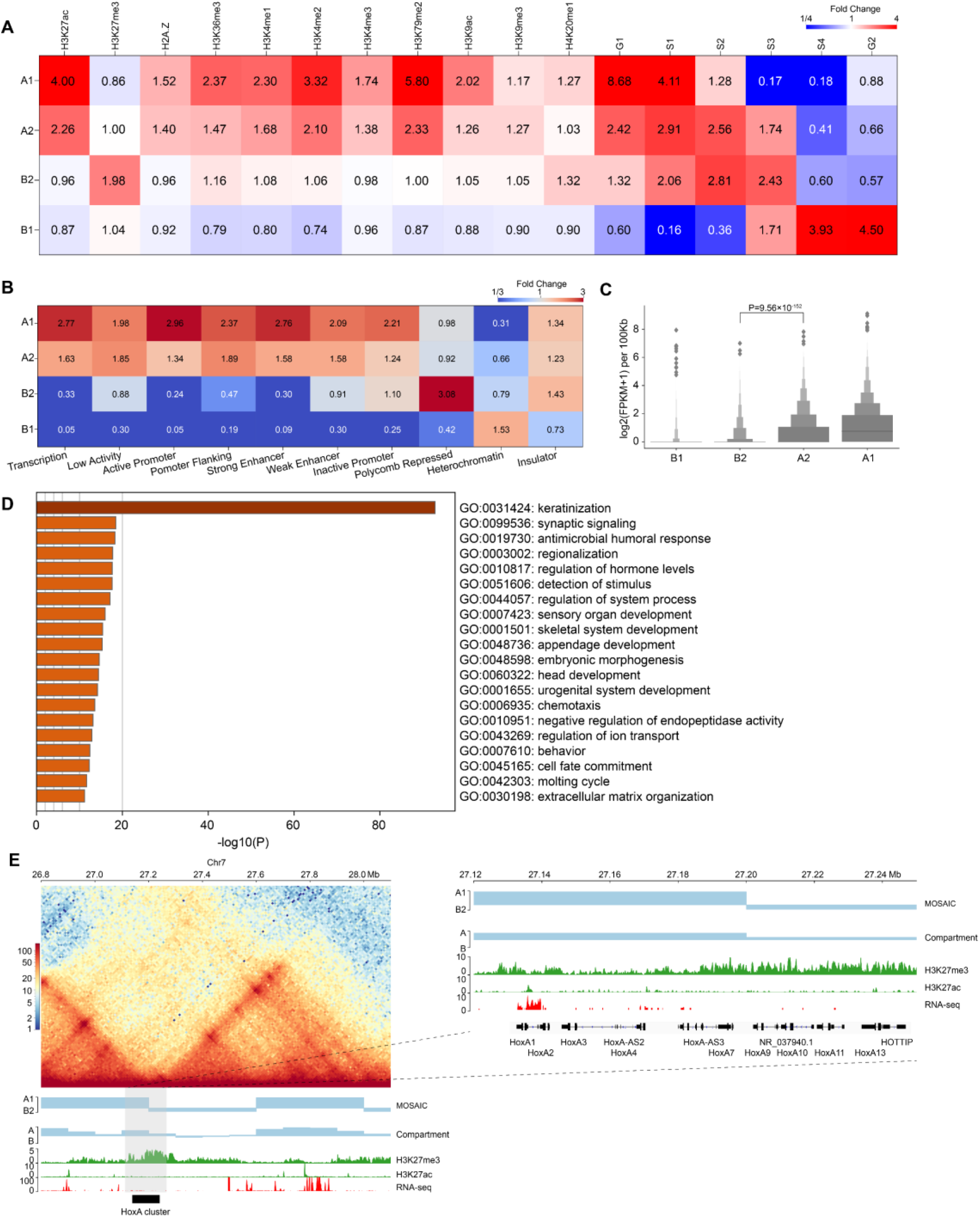
Compartmental states show distinct epigenomic features. (*A*) Heatmap of histone mark and replication timing enrichment for four compartmental states throughout the genome. (*B*) Heatmap of ChromHMM annotation enrichment for four compartmental states throughout the genome. (*C*) Gene expression of four compartmental states throughout the genome (P value calculated by one-tailed Mann–Whitney rank test). (*D*) Enriched GO terms of genes in B2. (*E*) Left panel: Hi-C interaction heatmap along with compartmental states, A/B compartment, H3K27me3, H3K27ac, and RNA-seq around the *HoxA* cluster. Right panel: zoom-in view of the region where the *HoxA* cluster is located.

In addition, we used chromatin state annotation by ChromHMM (Ernst and Kellis 2012) to examine their relation to compartmental states (Fig. 3B; Supplemental Fig. S2D). A1 is most enriched in the chromatin states “Transcription” and “Active Promoter”, and A2 is most enriched in “Low Activity” and “Promoter Flanking”. On the other side, B2 is highly enriched in “Polycomb Repressed” chromatin, and B1 is enriched in “Heterochromatin”. These results are consistent with the epigenetic pattern enrichment shown above, also support that functionally similar regions are in close spatial proximity (Hnisz et al. 2017).

We then compared gene expression among compartmental states (Fig. 3C). Consistent with the ChromHMM annotation results, A1 primarily contains expressed genes. Genes in A1 expressed at higher levels than A2. By contrast, B1 and B2 have minimal levels of transcription. Again, expression level of genes in A2 are significantly higher than those in B2, supporting our definition of the two types of compartments as euchromatin and heterochromatin, respectively. In summary, chromatin activities ascend in the order of B1, B2, A2, A1 in terms of gene expression. Then, we conducted Gene Ontology (GO) analyses using Metascape (Zhou et al. 2019). The result in Figure 3D shows that genes in B2 are enriched for developmental and cell type-specific functions. In particular, the late cornified envelope (LCE) gene cluster and keratin (KRT) gene cluster associated with keratinization are both located in B2 (Supplemental Fig. S3C). Overall, A2 and B2 are enriched in regulation-related genes (Supplemental Fig. S3A-B).

Considering the enrichment of regulation-related genes in A2 and B2, the compartmental states may contribute to their transcriptional regulation. Taking the genomic regions containing the *HoxA* cluster as an example, the corresponding EV1 values are positive, indicating that the whole *HoxA* cluster is in the conventional A compartment. In contrast, our scheme more accurately separated the cluster into A1 and B2, which is marked by H3K27me3 (Fig. 3E) (Lopez-Delisle et al. 2020). Our result is also more consistent with the finding that H3K27me3 system forms repressive microenvironments at *Hox* gene clusters in the more active nuclear environment in differentiated cells (Vieux-Rochas et al. 2015). This example highlights the necessity of recognizing regions as heterochromatic B2 embedded in large area of conventional euchromatic A compartment. Together, these results reveal that compartments identified by structural modularity are in different chromatin states and contain genes that perform different classes of functions in the cell.

### MOSAIC outperforms subcompartment identification methods in structural partition of individual chromosomes

Existing methods using inter-chromosomal contacts divide the genome into six subcompartments that are functionally and spatially distinct (Rao et al. 2014; Xiong and Ma 2019; Ashoor et al. 2020). It is of interest to compare the results of subcompartment identification using inter-chromosomal interactions and MOSAIC based on intra-chromosomal interactions. For this purpose, we used clustering metrics and modularity to quantify the effects of different compartmental partitions. We adopted the Silhouette coefficient (Rousseeuw 1987) and Calinski-Harabasz score (Calinski and Harabasz 1974) as indicators of consistency within a cluster. The Silhouette coefficient ranges from −1 to 1, whereas the Calinski-Harabasz score, as a standard of variance ratio, has values greater than zero. In both cases, higher values indicate better clustering performance. Modularity is a system property for characterizing community structure in networks, i.e., dense connection within groups but sparse connections between groups (Newman 2006). Modularity is utilized to measure the independency of clusters. Higher modularity indicates closer proximity of regions in the same cluster. The three available subcompartment identification methods are denoted as Rao_HMM (Rao et al. 2014), Xiong_SNIPER (Xiong and Ma 2019), and Ashoor_SCI (Ashoor et al. 2020), respectively. As shown in Figure 4A, MOSAIC outperforms all subcompartment identification methods in the division of individual chromosomes, in terms of both cluster consistency and independence.

**Figure 4.**
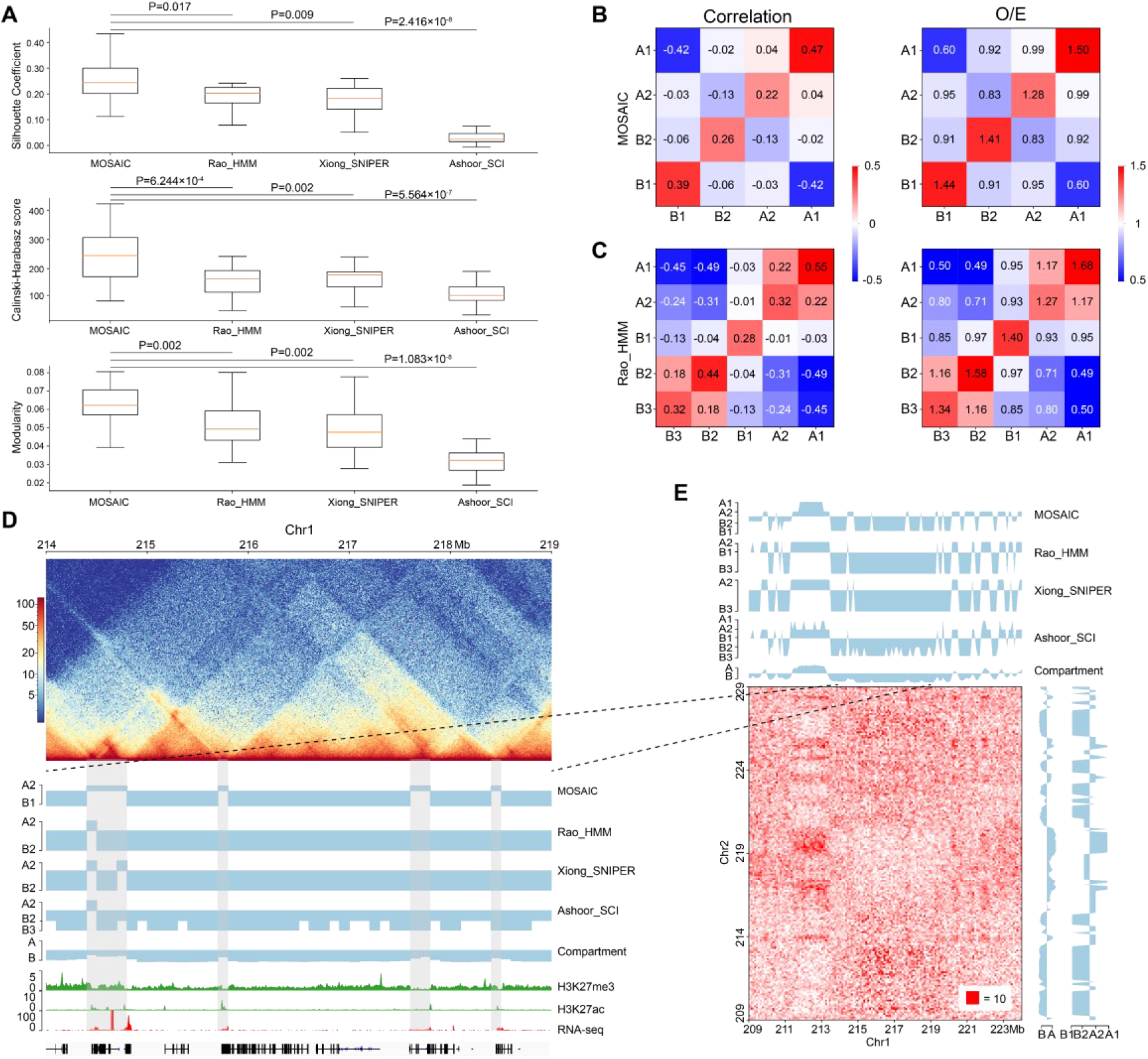
Comparisons of MOSAIC with the subcompartment identification methods. (*A*) Silhouette coefficients (top), Calinski–Harabasz scores (middle), and modularity (bottom) were used to compare the results of MOSAIC, Rao_HMM, Xiong_SNIPER and Ashoor_SCI in the division of individual chromosomes (P values calculated by one-tailed Mann–Whitney rank test). (*B-C*) Heatmap of mean values of the correlation matrix (left panel) and mean values of the O/E matrix (right panel) of various compartmental state combinations identified by MOSAIC (*B*) and Rao_HMM (*C*). Because subcompartment B4 was identified only on chromosome 19, it was excluded from this analysis. (*D*) An example showing that MOSAIC reveal more detailed and accurate compartmentalization at the intra-chromosomal level than subcompartments at the three shaded areas. (*E*) Inter-chromosomal heatmap showing that the specific compartmental states (shaded areas) identified by MOSAIC in (*D*) also have different interaction patterns relative to the surrounding regions.

While O/E matrix reflects direct interactions, the correlation matrix measures the relationships of interaction patterns. We utilized both types of information at the intra-chromosomal level to measure the clustering effect (Fig. 4B–C; Supplemental Fig. S4A-B). The compartmental states identified by MOSAIC consistently exhibit preferential interactions within the same states and depleted interactions between different states. The only exception is the marginal enrichment (0.04) between A1 and A2 in the correlation matrix. By contrast, for subcompartments of Rao_HMM, A1-A2 and B2-B3 exhibit enriched interactions. When we compared the extent of segregation between compartmental states, we observed the strongest segregation for A1-B1, followed by A2-B2. This is consistent with the results of the SVD analysis (Fig. 1G), in which A1 and B1 are separated along EV1, whereas A2 and B2 are separated along EV2. Given the strong depletion of A1-B1 and A2-B2 in terms of genomic neighborhood (Fig. 2D), these two pairs tend to segregate in both 1D and 3D.

Because subcompartments are obtained using genome-wide inter-chromosomal data, they may not consistently reflect fine interaction patterns within individual chromosomes. As shown in Figure 4D, there are four shaded areas, three of which are 100–200 kb regions not discerned by subcompartment identification methods. MOSAIC distinguishes all three as A2. Histone modification and gene expression information support the distinction of these regions from their surroundings as active chromatin. Moreover, the inter-chromatin interaction matrix shows that the MOSAIC-specific compartmental state A2 exhibits a different interaction pattern than the surrounding B1 region. We can also see that the MOSAIC-specific compartmental states A2 in chromosome 1 have weak interactions with A2 on chromosome 2. The reason why this A2 was not detected by subcompartment identification methods might be that the inter-chromosomal interactions are not as pronounced as the intra-chromosomal interactions (Fig. 4E). Taken together, MOSAIC exhibits higher accuracy and sensitivity than subcompartment identification methods in characterizing chromatin compartmentalization at intra-chromosomal level.

### Compartmental states accurately reflect gene expression dynamics

Since GM12878 and K562 are closely related, we compared these two cell lines to investigate the relationship between compartmental state and gene expression. We identified the regions in which compartmental states change and then analyzed differentially expressed genes (DEGs). Among the 1,420 DEGs (>50% change in FPKM and P value < 0.05) (Djebali et al. 2012), 533 are in regions with compartmental changes using MOSAIC. By comparison, only 234 DEGs are in regions with switches in A/B compartments. Among the 533 DEGs, 194 are shared between the two methods, whereas the other 339 are MOSAIC-specific (Fig. 5A; Supplemental Fig. S5A).

**Figure 5.**
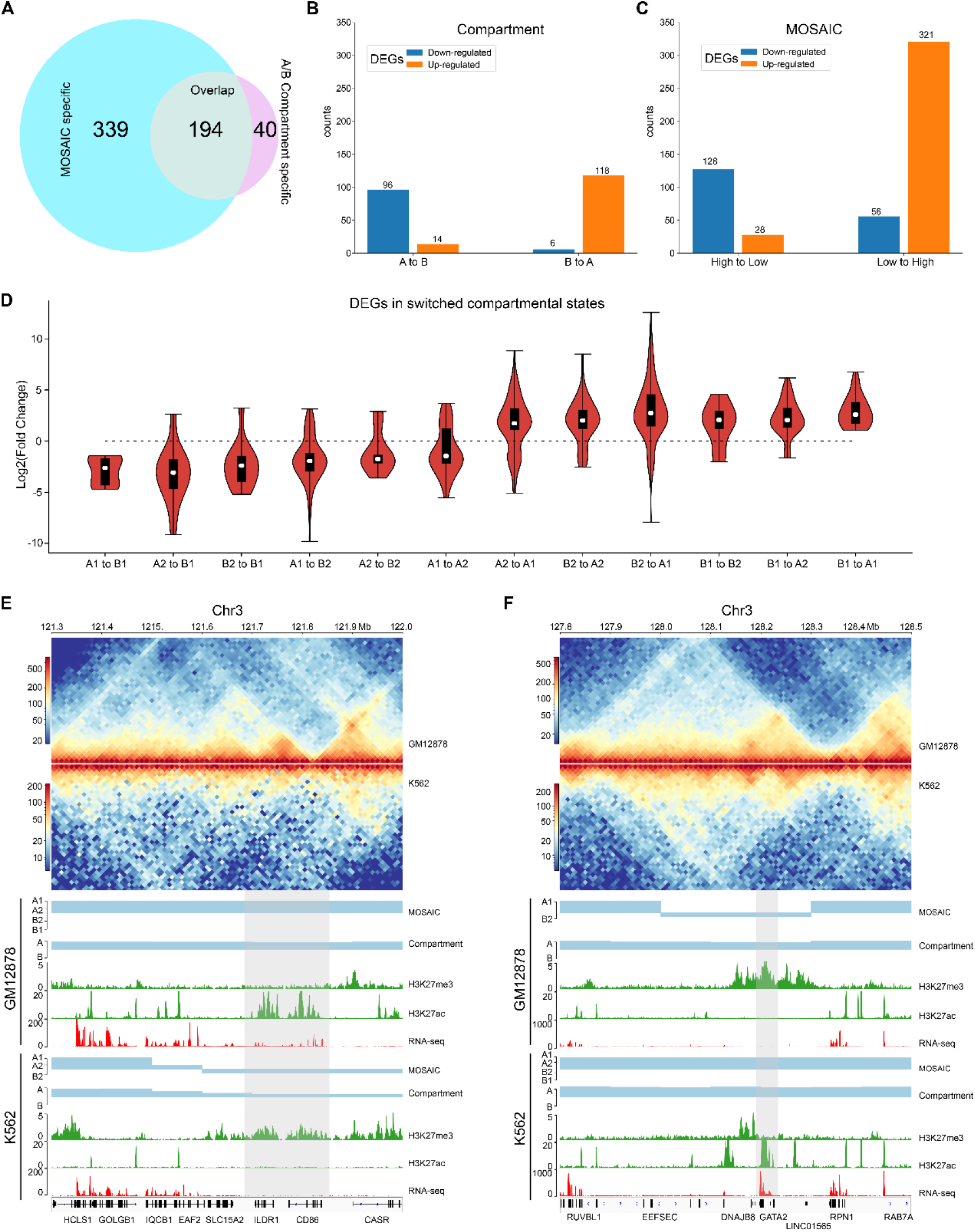
MOSAIC reveals concordance of gene expression and compartmental state dynamics between K562 and GM12878 cells. (*A*) Venn diagram showing the overlap of DEGs that can be covered by the two schemes of compartmentalization analysis. (*B-C*) Boxplots showing categorization of DEGs with A/B compartment switches (*B*) and with compartmental state switches (*C*). For DEGs, the ratio of gene expression level in GM12878 to K562 were calculated. Switches in compartmental state are grouped into descending and ascending order, in terms of transcription activity implications of compartmental states. (*D*) Fold change distribution of DEGs grouped by compartmental state switches. Fold changes were calculated as *log*_2_(*FPKM*_*GM*12878_/*FPKM*_*K*562_). (*E-F*) Two examples on chromosome 3 showing concordance between compartmental states identified by MOSAIC and expression of *CD86*, *ILDR1* (*E*) and *GATA2* (*F*) located in the shaded areas. In both cases, A/B compartment scheme does not identify switch of compartment.

Next, we examined the extent of concordance between gene expression and compartmental states. Figure 5B shows that only 214 DEGs are consistent with A/B compartment switches. By contrast, MOSAIC identified 449 DEGs that exhibit concordance between compartmental states and gene expression (Fig. 5C). Particularly, as shown in Figure 5D and Supplemental Figure S5B, compartmental changes are accompanied by consistent changes in gene expression: genomic regions changing toward more active compartments contain genes that are upregulated, and vice versa. This suggests that MOSAIC scheme is much more sensitive than conventional A/B compartment scheme in characterizing the dynamics of transcriptional regulation. A/B compartment scheme might severely under-estimate the chromatin architectural basis of transcriptional regulation.

Figure 5E and 5F show two typical regions containing DEGs without A/B compartment switches. In the A/B compartment scheme, the regions shown in Figure 5E are constantly A compartment in both GM12878 and K562. However, RNA-seq data showed that two genes, *CD86* and *ILDR1*, were specifically expressed in GM12878. This differential expression was accompanied by dynamic H3K27ac and H3K27me3 signals. MOSAIC reconciles the dynamics by revealing that this region switches from A1 in GM12878 to B2 in K562. Interestingly, both *CD86* and *ILDR1* are associated with immunoglobulin, which is related to the function of GM12878 cells. A recent study on GM12878 cells showed that this region contains a super-enhancer that regulates both genes (Kleinstern et al. 2020). Moreover, a genome-wide association study validated one locus at 3q13.33 (rs9831894) that is significantly inversely associated with the risk of diffuse large B-cell lymphoma (DLBCL) (Kleinstern et al. 2020). Accurate compartmentalization characterization by MOSAIC might provide insight of this phenotypical difference from the perspective of fine-scale chromatin architecture dynamics.

As shown in Figure 5F, the entire region belongs to the A compartment in both K562 and GM12878 cells according to EV1 values. By contrast, MOSAIC identified the region containing *GATA2* as B2 in GM12878 and A1 in K562. *GATA2* encodes a transcription factor that is involved in stem cell maintenance and plays a significant role in hematopoietic development (de Pater et al. 2013). From a functional standpoint, *GATA2* should be specifically expressed in K562. Consistent with this, in GM12878 cells, the genomic region containing *GATA2* has an elevated level of H3K27me3 to repress its expression.

In summary, the capacity to interpret differential expression between cells based on changes in the A/B compartment was limited. By contrast, concordance between gene expression and compartmentalization was significantly improved in MOSAIC. The fine-scale compartmental dynamics involving micro-compartments revealed by MOSAIC might shed light on mechanisms of transcriptional regulation of genes important for cell-type specific function and disease.

## Discussion

We applied SVD to intra-chromosomal contacts by extending the conventional A/B bipartition which uses only EV1 to a comprehensive compartmentalization framework using further EVs. Although EV1 matches the overall plaid patterns, EV2 provides a striking separation of ambiguous regions with marginal EV1 values, also highlighted by corresponding bimodality of H3K27me3. The regions pinpointed by EV2 are typically interspersed with activities opposite to their surrounding environments. Specifically, A2 and B2, referred to as micro-compartments, are embedded in large areas of B1 and A1, respectively. Our scheme provides a fine-scale and dynamic view of chromatin compartmentalization. While EV1 characterizes the euchromatin (A1) and constitutive heterochromatin (B1) corresponding to the structurally stable parts of the chromatin, bipolar EV2 captures the dynamic aspects of chromatin that are structurally flexible and functionally regulated. Figure 6 gives a schematic representation of fine-scale chromatin compartmentalization and its dynamics in the space spanned by the top two EVs.

**Figure 6.**
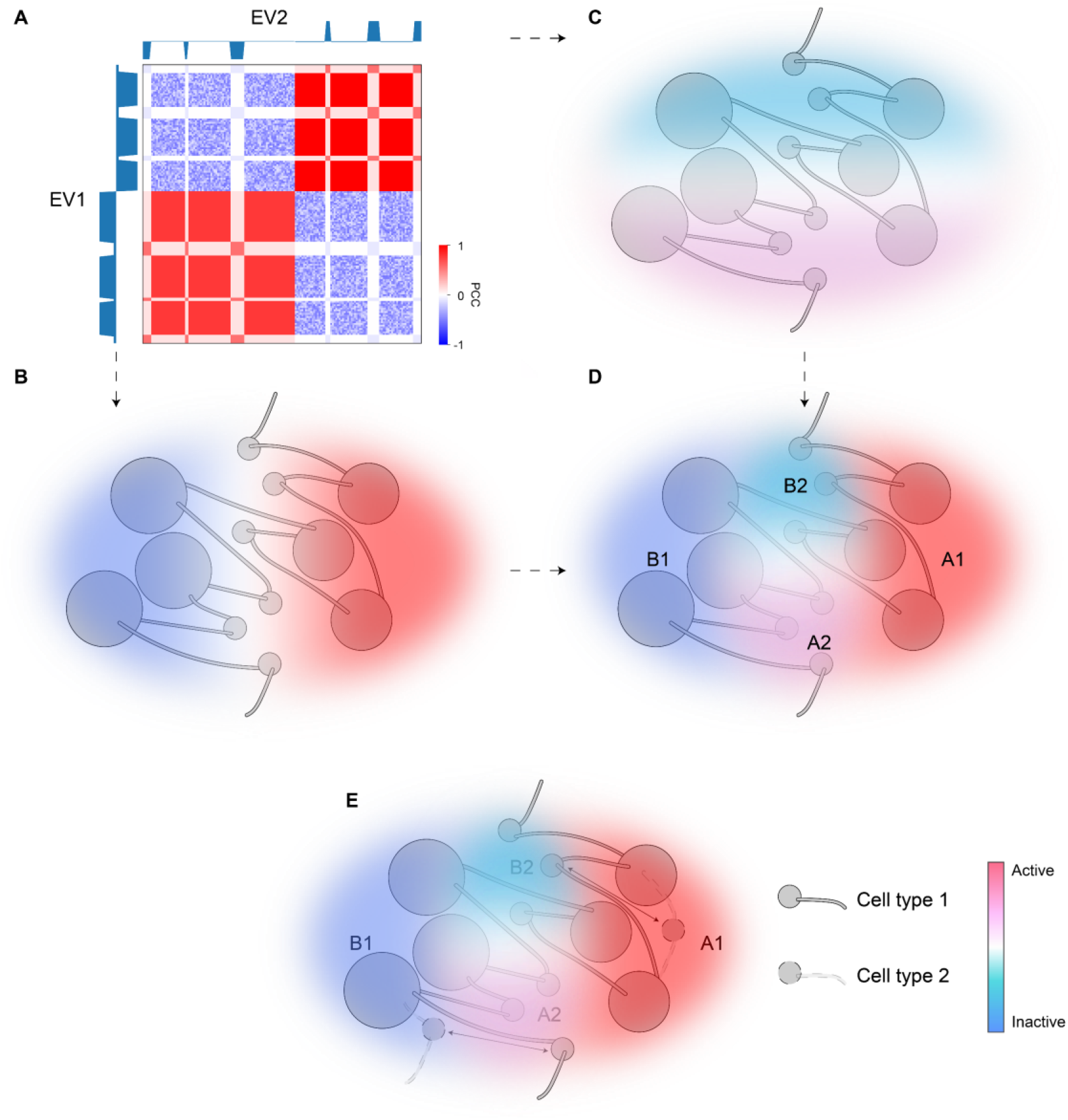
A model of dynamic chromatin compartmentalization highlighting genomic and architectural features of micro-compartments. (*A*) The schematic Hi-C heatmap of an area of the genome containing A1, A2, B1, and B2 compartment, identified based on EV1 and EV2. EV1 can separate this segment into two major categories (B1, left; A1, right), but cannot distinguish between the small regions (A2/B2) embedded in each major category, whereas EV2 marks A2 and B2. (*B*) The sign of EV1 can separate A1 and B1. Color represents activities (red for active; blue for inactive). (*C*) The sign of EV2 can separate A2 and B2. Color represents activities (red for active; blue for inactive). (*D*) The combination of EV1 and EV2 can accurately identify all four compartmental states and accurately reflect their activities. (*E*) A diagram showing the dynamic regions between the two cell types are the micro-compartment A2 and B2. Arrows indicate cell-type specific compartmental states for certain regions.

In comparison with other state-of-the-art compartment identification tools (Durand et al. 2016; Wolff et al. 2018) based on PCA or SVD, MOSAIC provides automatic and accurate facilities allowing universal identification of potentially diverse compartmentalization patterns. It is recognized that the first PC in PCA may not represent compartmental features for all chromosomes (Schmitt et al. 2016b). Consequently, manual curation is required for proper PC selection in current A/B compartments identification tools. Using modularity as the criterion, we demonstrated the power of MOSAIC for automatically selecting EVs that correctly reflect structural and functional features of chromatin. MOSAIC scheme is also robust in the sense that it accommodates scenarios with structure-function relations that might differ among diverse species and cell types. For instance, the layout of some chromosomes may have more than four compartmental states, or have histone modifications other than H3K27me3, which is indicative of its fine-scale structure. MOSAIC allows detection of such compartmental structures by proper, automatic EV selection and robust clustering. Considering the A/B compartments based on the leading EV as a first-order approximation of the true compartmental states, MOSAIC represents a natural extension that uncovers finer-scale patterns by including the second EV. MOSAIC framework also allows generalization to EVs that further qualify the structural and functional analysis described above.

Currently, conventional A/B compartment scheme is widely adopted for comparative analysis on chromatin compartmentalization. However, comparison between GM12878 and K562 cells using MOSAIC revealed extensive and fine-scale concordance between compartmental state and gene expression which are missed by A/B compartments. The rationale behind this finding is that transcriptional regulation through switching between constitutive euchromatin and heterochromatin is rarer than expected. Instead, our results show that the majority of compartmental neighborhood and dynamics are A1-B2 and B1-A2. Incapability of A/B compartment scheme in capturing such types of compartmental neighborhood and dynamics leads to its limitations. Consequently, the analysis based on bipartite compartments is prone to under-estimate the role of compartmentalization in transcriptional regulation, exemplified by the cases of *CD86*, *ILDR1* and *GATA2* shown above.

Spatial segregation of regions marked by H3K27me3 has been observed in various systems and loci (Vieux-Rochas et al. 2015; Du et al. 2020; Johnstone et al. 2020; Heurteau et al. 2020). For example, a study focusing on *Hox* gene clusters revealed clustering of these regions with other H3K27me3 targets (Vieux-Rochas et al. 2015). Another investigation of mouse development revealed Polycomb-associating domains and local compartment-like structures (Du et al. 2020). In addition, a study (Johnstone et al. 2020) comparing primary and tumor cells found an intermediate compartment highlighted by DNA hypomethylation and H3K27me3. Based on this observation, the authors proposed a three-compartment model. Generalizing the above findings, our work provides a global and comprehensive view of the role of H3K27me3 in compartmentalization within the framework of MOSAIC. In each chromosome, EV2 correlates with H3K27me3. The strong modularity of the resultant B2 compartment supports the aggregation of the H3K27me3-marked regions. Furthermore, our results show that the typical H3K27me3-marked regions are embedded in large areas of the conventional A compartment, consistent with the contrasting active nuclear environment of repressive *Hox* gene clusters previous observed (Vieux-Rochas et al. 2015).

A recent study (Kundu et al. 2017) based on 5C using ESCs revealed PRC1-mediated self-interacting domains that are similar to TADs but smaller in size. Our observations of *HoxA* cluster in GM12878 cells suggest specific compartmental states corresponding to H3K27me3-marked regions (Fig. 3E). While these results support the role of H3K27me3 in chromatin architecture at various layers, the interplay between domain and compartmental states, and the underlying histone modifications, remains to be elucidated. Based on the close connection between H3K27me3 and micro- compartments, it is promising that compartmentalization might play a significant role in histone mark spreading.

While TADs represent local segregation of adjacent genomic regions, compartments represent global aggregation of 1D-separated regions with similar genomic features and epigenetic and functional states. Under the current working model, TADs influence transcription by providing a local environment that facilitates enhancer-promoter looping within the same TAD and insulates enhancer-promoter contact across TADs (Andrey et al. 2013). However, the perturbation experiments (Schwarzer et al. 2017; Rao et al. 2017) removing TADs and loops did not lead to proportionally differential gene expression. It is important to note that these perturbations reinforced compartmentalization, pointing to the potential role of compartmentalization in transcriptional regulation, which might be under-estimated by conventional comparative analysis based on A/B compartments as shown above. The extensively improved concordance between expression and compartmental states revealed by MOSAIC supports transcriptional regulation through compartmentalization as an important mechanism.

Hierarchical modularity is a general organizing principle of living systems. From the perspective of hierarchical chromatin architecture in the nucleus (Gibcus and Dekker 2013), compartmental states add another layer to hierarchy of its spatial modular organization. In this hierarchy, the well-known chromosome territories represent the highest level of modularity, highlighted by chromosome-specific radial distribution within the nucleus. Under this top-level circumstance, A/B compartments form the second level of organization which spatially segregate chromatin with different activities at a large scale. Embedded in large areas of A or B compartments, fine-scale regulation of individual genes toward the opposite activities, represent the third level of architecture, in which loci are spatially segregated into micro-compartments. The precise and delicate regulation of genome function necessitates this hierarchical compartmentalization of chromatin architecture. Corresponding perturbation experiments might provide exact answer to the mechanistic connection between chromatin architecture and function.

## Methods

### A/B compartment identification

Compartment A and B are determined by the sign of the first eigenvector obtained from PCA of intra-chromosomal Hi-C contact maps. Regions with opposite EV1 signs in different samples are considered as compartment switching between A and B.

### The MOSAIC algorithm

#### Step 1: Remove centromere effect and matrix reconstruction

Because the O/E matrix is a symmetric and non-negative adjacency matrix of the chromosomal contact map, SVD on O/E matrix *M* can get the eigenvalues and eigenvectors according to the following equation:

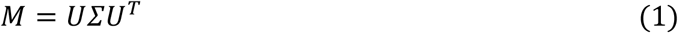

where

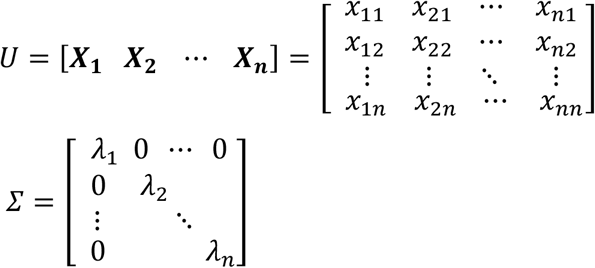

Here *U* is a matrix composed of eigenvectors, and *Σ* contains the eigenvalues in descending order of magnitude.

The top eigenvectors are of particular importance because they reflect the dominant structure of the O/E matrix. In general, the top eigenvectors represent genome coverage, global or local pattern, centromere effect, etc. The eigenvector with strong bias between chromatin’s p and q arm is considered to represent the centromere effect. Here, we define *I* to measure the effect using the following equation:

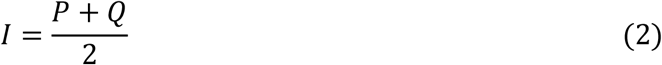

where

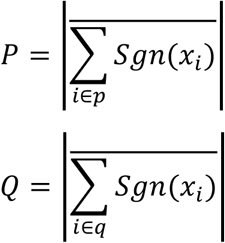

Here *x*_*i*_ is the value of the eigenvector at position *i*, and *p* and *q* represent the short and long arm, respectively. The *Sgn* function returns the sign of a number.

*I* ranges from 0 to 1. The higher the value of *I*, the stronger the arm effect. To disentangle the influence of arm effect from the compartmentalization pattern, the O/E matrix *M* was updated by modified eigenvectors and eigenvalues according to following equation:

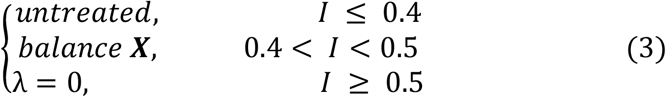

When *I* is ≥ 0.5, we replace the eigenvalue corresponding to the eigenvector with 0. For *I* between 0.4 and 0.5, we balance the corresponding eigenvector of the long arm or the short arm by subtracting their arm-wise means. We reconstruct the new O/E matrix *A* using the processed eigenvectors and eigenvalues. Without the above correction on centromere effect, the results will have negligible effect for most chromosomes. For the other several chromosomes with particularly strong centromere effect, the identified A2 or B2 will be restricted to within the long or short arms of the chromosomes, respectively, which is of course does not reflect the compartmental pattern.

#### Step 2: Pick eigenvectors with the highest and second-highest modularity

We perform SVD on the updated O/E matrix *A* to obtain the new eigenvectors. We then construct a Hi-C interaction graph by *A* and compute the modularity matrix, *B*, according to the following equation (Newman 2006):

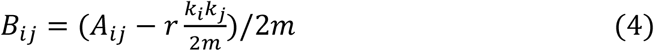

where *A*_*ij*_ is the new O/E matrix, *k*_*i*_ is the degree of *i*, *m* is total number of links, and *r* is the resolution parameter (*r* = 1 is used to compute the modularity). Thus, we can obtain the modularity *Q* of the whole network from modularity matrix *B* according to the following equation:

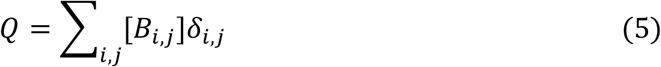

where *δ*_*i,j*_ is 1 if vertex *i* and vertex *j* are assigned to the same community, and 0 otherwise.

For each eigenvector, we can divide the chromosome corresponding to matrix *A* into two communities according to the signs of the values, so that we can calculate the modularity corresponding to each eigenvector. We take top two eigenvectors in terms of modularity for the next step of the analysis.

#### Step 3: K-means clustering

We perform K-means clustering on EV1 and EV2 weighted by eigenvalues by setting *k* equal to 3, 4, 7, or 8 according to SSE with *k* of SSE (Supplemental Fig. S1G). The best results were obtained when *k* was set to 4, and these results were consistent overall among chromosomes.

#### Step 4: Refine clustering by Louvain algorithm

To optimize the clustering results, we employed the Louvain algorithm to maximize modularity. The Louvain algorithm relies on an iterative method to quickly converge to the maximum in *Q*. For each iteration *t*, individual node movement maximizes the increase in modularity *Q*, Δ*Q*, according to the following equation:

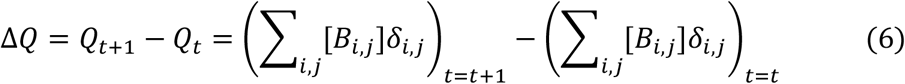

Initially, each node is assigned to the community according to the result of the K-means clustering. There are two ways to start the iteration: the reassignment process of the nodes in the first iteration is random, or the reassignment process of the nodes in the second iteration follows the order of their positions along the chromosome. We can choose any one of these two approaches for iteration and repeat until a local maximum of modularity is reached. The choice of approach has minor impact on the results. But to keep the results consistent, we used the second option.

### Data preprocessing of RNA-seq

We used StringTie (Pertea et al. 2015) to obtain the FPKM (Fragments per KB per million mapped reads) of each gene of GM12878 and K562, and then used Ballgown (Frazee et al. 2015) to identify DEGs between the two cell lines.

### Data preprocessing of epigenomic tracks

We use bwtool (Pohl and Beato 2014) to calculate the mean value of the histone and replication time signals in each 100 kb or 10 kb bin of downloaded bigWig tracks (see data availability). For the histone signal analysis in Fig. 1E and 1F, we adopt the Z-score to normalize the signal values.

### Enrichment analysis of epigenomic signal and chromatin state

For analysis of epigenomic signal enrichment and chromatin state enrichment, we adapted a previously described method (Rao et al. 2014). Enrichment of the epigenomic signal can be calculated as the median within the cluster divided by the total median at a resolution of 100 kb. For chromatin state enrichment calculations, we first binned the annotations into 100 kb bins and then calculated the proportion of each annotation within each bin. Finally, we divide the mean value of the proportion within the cluster by the mean value of the proportion across all bins. To facilitate the display of annotations in Fig. 3B, we abbreviate the annotations defined by ChromHMM, replacing “Transcription Associated” with “Transcription”, “Low activity proximal to active states” with “Low Activity”, “Candidate Strong Enhancer” with “Strong Enhancer”, “Candidate Weak enhancer/DNase” with “Weak Enhancer”, “Heterochromatin/Repetitive/Copy Number Variation” with “Heterochromatin” and “Distal CTCF/Candidate Insulator” with “Insulator”, respectively.

### Compartment border evaluation

Nine histone modification marks representing euchromatin, heterochromatin, and Polycomb-repressed chromatin were used to evaluate the effect of insulation between compartmental states. Briefly, we divided 10 bins on the left and right side of each compartment border and recorded the mean value of the histone modification signal in each 10 kb bin. Then, a N*20 matrix was generated in which the 20 columns represent the signal strength of two continuous compartments, and the N rows represent borders.

### GO analysis

Because A1 and A2 represent euchromatin, whereas B1 and B2 represent heterochromatin and Polycomb-repressed chromatin, respectively, we performed GO analysis on the transcribed protein-coding genes in A1 and A2, and the non-transcribed protein-coding genes in B1 and B2. B1 and B2 contain 1,385 and 1,335 non-transcribed protein-coding genes, respectively, whereas A1 and A2 contain 7,605 and 1,667 transcribed protein-coding genes, respectively. Because common GO analysis software, including Metascape, limits the number of input genes to 3,000 or fewer, we randomly selected 3,000 genes from A1 as input to Metascape to search for enriched GO terms.

### Compartmentalization strength

We obtained the corresponding eigenvectors after SVD of the O/E matrix *M*. Each eigenvector can separate the O/E matrix *M* into two parts through its sign. We define the compartmentalization strength of each eigenvector based on the compartmentalization (Schwarzer et al. 2017) defined as:

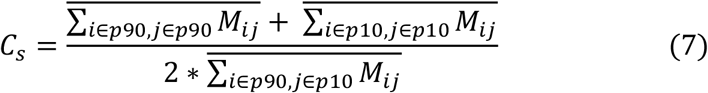

where *p*90 represents the location of the values above the 90^th^ percentile in this eigenvector, and *p*10 represents the location of the values below the 10^th^ percentile.

### Clustering metrics

The Silhouette coefficient is defined as:

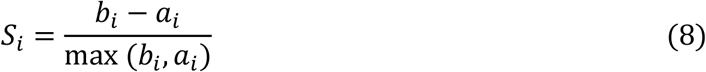

where *b*_*i*_ is the mean nearest-cluster distance for all other clusters, and *a*_*i*_ is the mean intra-cluster distance for genomic bin *i*. We calculated the Silhouette coefficient for each genomic bin; the mean Silhouette coefficient of all genomic bins was used to evaluate the quality of the clustering. Cosine distance was used as our distance measure.

For a set of data *E* that has been clustered into *k* clusters of size *n*_*E*_, the Calinski– Harabasz score is defined as the ratio of the sum of inter-cluster dispersion and intra-cluster dispersion for all clusters:

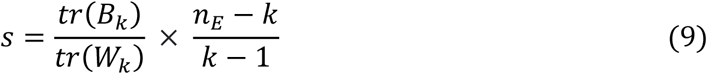

where *tr*(*B*_*k*_) is the trace of the inter-cluster dispersion matrix, and *tr*(*W*_*k*_) is the trace of the intra-cluster dispersion matrix, defined by:

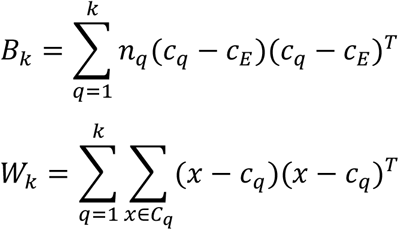

where *C*_*q*_ is the set of points in cluster *q*, *c*_*q*_ is the center of cluster *q*, *c*_*E*_ is the center of *E*, and *n*_*q*_ is the number of points in cluster *q*.

We used the silhouette_score and calinski_harabaz_score functions from scikit-learn package version 0.19.0 for the calculations.

### Data access

All downloaded data are based on the hg19 reference genome.

Compartmental conservations were obtained from 5 cell lines and 11 tissues.

The Hi-C data of 5 cell lines (GM12878, K562, IMR90, NHEK, HUVEC) were obtained from ftp://cooler.csail.mit.edu/coolers/hg19/ in cool format but original data were from Rao et al. in GEO accession GSE63525.

The Hi-C data of 11 tissues (Adrenal, Bladder, Dorsolateral Prefrontal Cortex, Hippocampus, Lung, Ovary, Pancreas, Psoas, Right Ventricle, Small Bowel, Spleen) were obtained from GSE87112.

The signal bigWig tracks of GM12878 for histone modifications were obtained from the ENCODE consortium using the following link: https://www.encodeproject.org/.

The signal bigWig tracks of GM12878 and K562 for replication time data were obtained from the ENCODE consortium using the following link: http://hgdownload.cse.ucsc.edu/goldenPath/hg19/encodeDCC/wgEncodeUwRepliSeq/.

The RNA-seq data of GM12878, K562 in bam format were obtained from Djebali et al. in GEO accession number GSE33480.

The ChromHMM annotations of GM12878 in hg19 were obtained from ENCODE consortium using the following link: https://genome.ucsc.edu/cgi-bin/hgFileUi?db=hg19&g=wgEncodeAwgSegmentation.

The subcompartments identification of Rao_HMM was obtained from https://www.ncbi.nlm.nih.gov/geo/download/?acc=GSE63525&format=file&file=GSE63525%5FGM12878%5Fsubcompartments%2Ebed%2Egz.

The subcompartments identification of Xiong_SNIPER was obtained from https://cmu.box.com/s/n4jh3utmitzl88264s8bzsfcjhqnhaa0.

The subcompartments identification of Ashoor_SCI was obtained from https://github.com/TheJacksonLaboratory/sci/tree/master/predictions.

MOSAIC code is available on https://github.com/WenZi0809/MOSAIC.

## Acknowledgements

The authors would like to thank Victor Corces for insightful comments on the manuscripts. We acknowledge financial support from the National Natural Science Foundation of China (no. 31771430 to L.L., no. 31571347 to C.H.), Huazhong Agricultural University Scientific and Technological Self-innovation Foundation (to L.L.), Hubei Hongshan Laboratory (to L.L.) and Shenzhen Science and Technology Innovation Commission (no. JCYJ20170412152835439 to C.H.).

## Author’s contributions

L.L. conceived the project. Z.W. and L.L. designed the algorithm. Z.W. implemented the algorithms, developed software, and performed computational analysis. W.Z., Q.Z. and J.X. contribute to Hi-C data collection and pre-processing. C.H. and Z.Q. advised on data analysis. L.L. and Z.W. wrote the manuscript with input from all authors.

