## Supplemental Figures for "Extensive Chromatin Structure-Function Association Revealed by Accurate Compartmentalization Characterization"

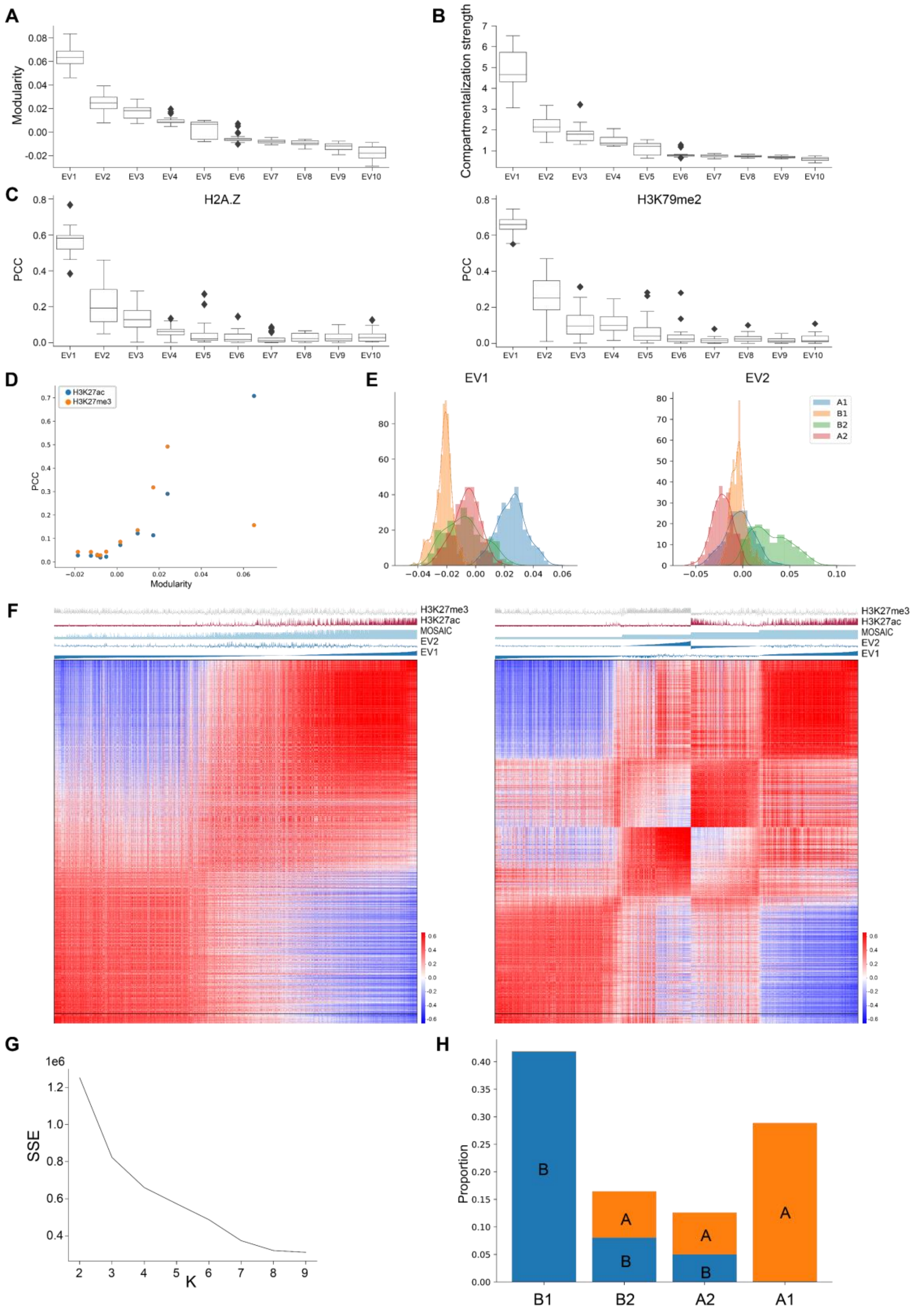


**Figure S1.** Exploration of EV1 and EV2. (*A*-*B*) Distribution of modularity (A) and compartmentalization strength (*B*) in the top-ten EVs, sorted by modularity. (*C*) Pearson correlation coefficients between H2A.Z (left) and H3K79me2 (right) and the top-ten EVs sorted by modularity. (*D*) Scatter plot of the average Pearson correlation coefficients to histone modifications of all chromosomes versus average modularity of all chromosomes for the top-ten EVs in each chromosome, sorted by modularity. (*E*) Distribution of A1, A2, B2, and B1 on EV1 (left) and EV2 (right) in chromosome 1. (*F*) Results of the segmentation of chromosome 1 with only EV1 (left) versus the combination of combining EV1 and EV2 (right). (*G*) Sum of squared errors of results obtained by K-means clustering of EV1 and EV2 with different K. (*H*) Proportion of areas of A1, A2, B2, and B1 in the A/B compartment scenario.

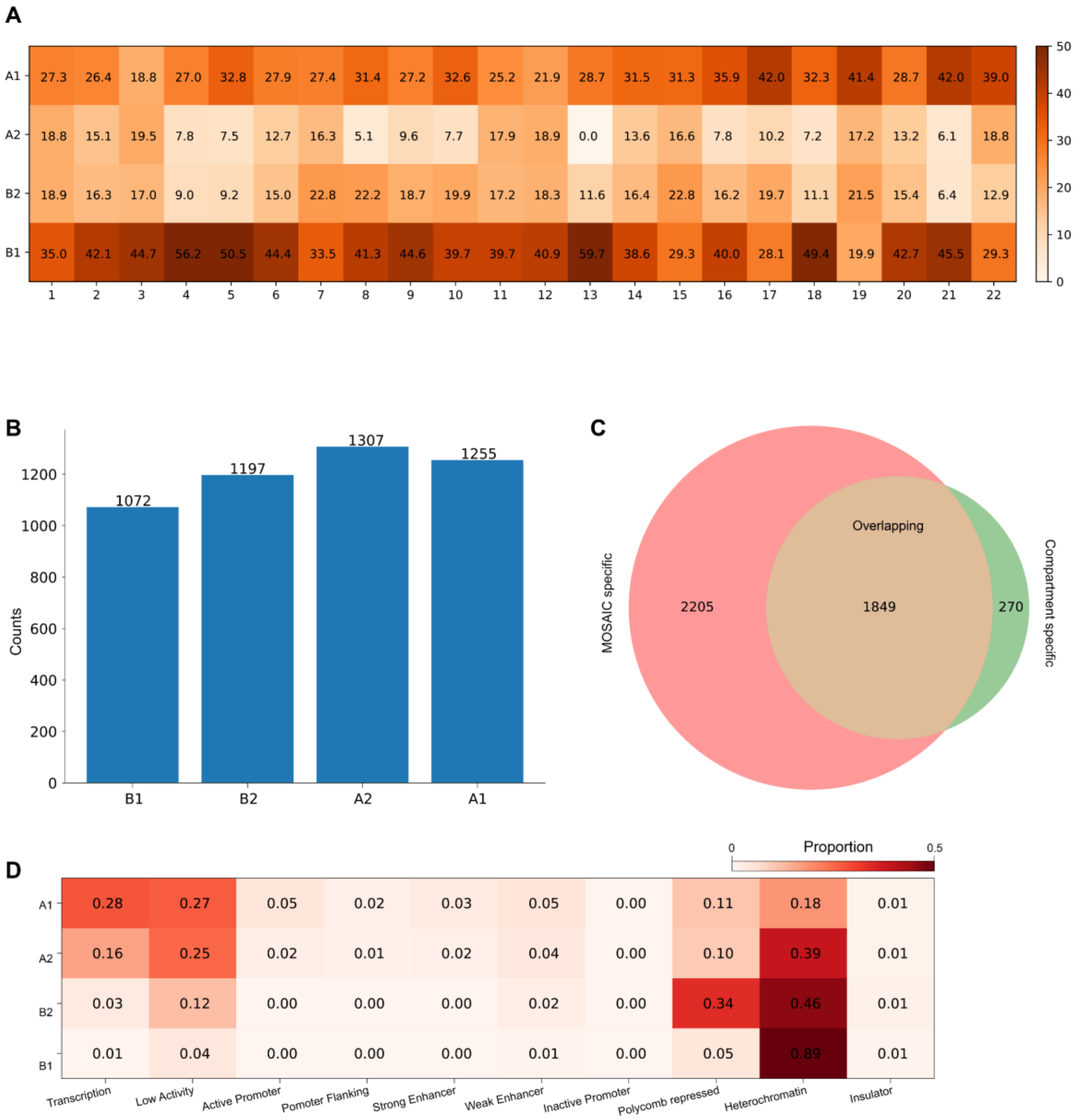


**Figure S2.** Characteristics of A2 and B2. (*A*) Cluster percentage for each chromosome. (*B*) The number of regions in each cluster. (*C*) The number of borders obtained by MOSAIC and A/B compartment scenario. (*D*) Heatmap of ChromHMM annotation proportion for four compartmental states throughout the genome.

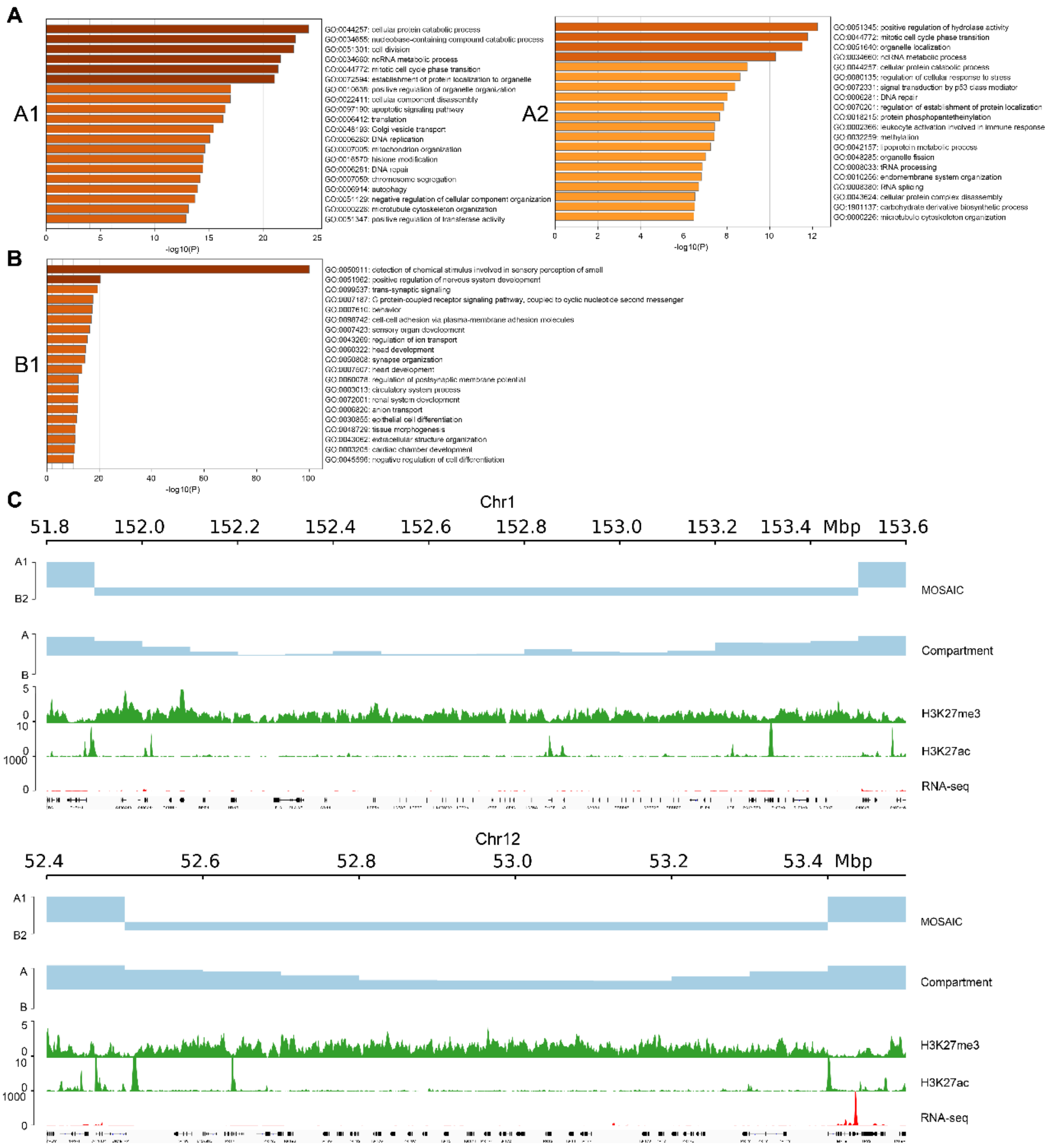


**Figure S3.** GO analysis of A1, A2, and B1. (*A*) Enriched GO terms in A1 (left) and A2 (right). (*B*) Enriched GO terms in B1. (*C*) LCE gene cluster (top) and KRT gene cluster (bottom) are all located in the B2 region identified by MOSAIC, but in the A region identified by A/B compartment scenario.

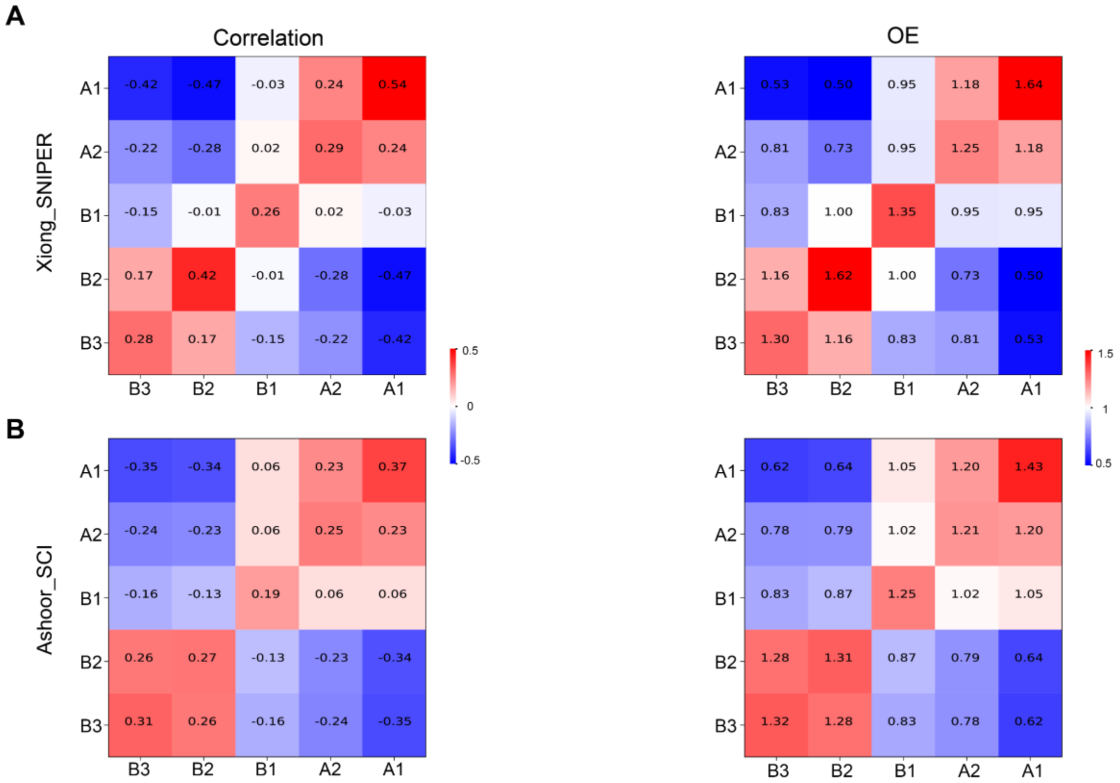


**Figure S4.** Evaluation of Xiong_SNIPER and Ashoor_SCI. (*A*-*B*) Heatmap of mean value of the correlation matrix (left panel) and mean value of the O/E matrix (right panel) in each compartmental state identified by Xiong_SNIPER (*A*) and Ashoor_SCI (*B*).

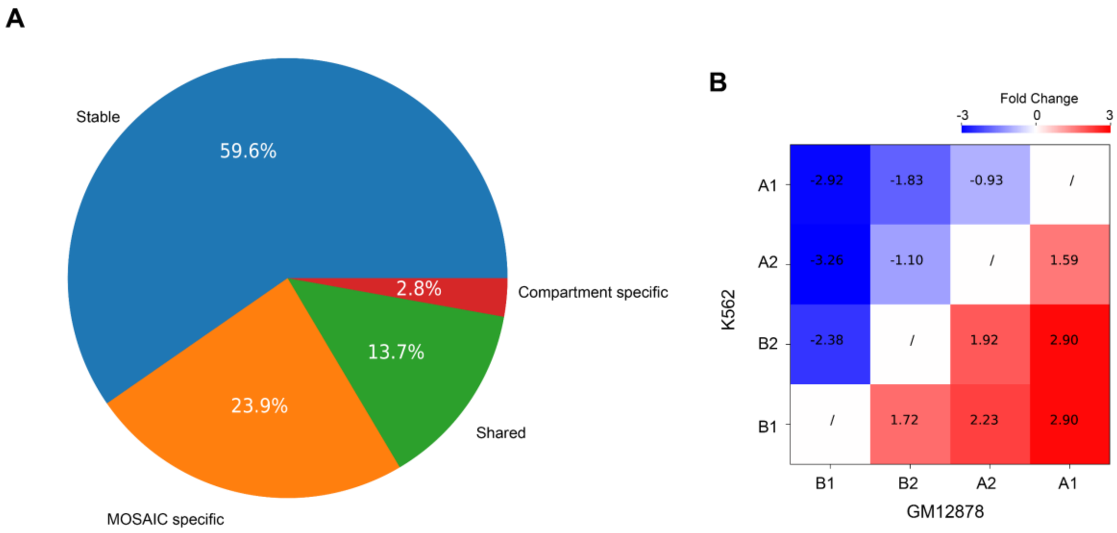


**Figure S5.** Percentage of DEGs between GM12878 and K562 in the MOSAIC scenario and the magnitude of the change. (*A*) Percentage of DEGs covered by MOSAIC and A/B compartment scenario. (*B*) The mean value of fold changes in DEGs in switches between compartmental states.
